# CoPISA: Combinatorial Proteome Integral Solubility/Stability Alteration analysis

**DOI:** 10.64898/2026.03.20.713131

**Authors:** Ehsan Zangene, Elham Gholizadeh, Uladzislau Vadadokhau, Danilo Ritz, Amir A. Saei, Mohieddin Jafari

**Affiliations:** Department of Biochemistry and Developmental Biology, Faculty of Medicine, University of Helsinki, Helsinki, Finland; Biozentrum, University of Basel, Basel, Switzerland; Department of Microbiology, Tumor and Cell Biology, Karolinska Institutet, Stockholm, Sweden; Faculty of Medicine and Health Technology, Tampere University and TAYS Cancer Center, Tampere, Finland; Tampere Institute for Advanced Study, Tampere University, Tampere, Finland

## Abstract

Combination therapies are widely used in acute myeloid leukemia (AML), but systematic datasets capturing proteome-wide responses to multi-drug perturbations remain limited. Here we present CoPISA (Combinatorial Proteome Integral Solubility/Stability Alteration), a quantitative proteomics assay designed to profile protein solubility and stability responses to single and combined drug treatments. The dataset includes two AML drug pairs (LY3009120–sapanisertib and ruxolitinib–ulixertinib) applied to four AML cell lines (MOLM-13, MOLM-16, SKM-1, and NOMO-1) under control, single-agent, and combination conditions in both lysate and intact-cell formats. Thermal solubility profiling coupled with TMT-based multiplexed LC–MS/MS generated 16 TMT16-plex experiments comprising 192 LC–MS/MS raw files, providing deep proteome coverage across treatments and biological contexts.

The resource includes raw and processed proteomics data, detailed experimental metadata in Sample and Data Relationship Format (SDRF), and reproducible analysis scripts for reporter normalization, protein-level aggregation, statistical modeling, and classification of combinatorial response patterns. The experimental design enables identification of proteins responding uniquely to combination treatments as well as overlapping single-agent effects. Technical validation demonstrates reproducible quantification across multiplex experiments and assay formats. All data are publicly available through the PRIDE repository (PXD066812) together with analysis code, enabling independent reanalysis and method development. This dataset provides a benchmark resource for studying proteome responses to drug combinations, comparing lysate and intact-cell perturbation profiles, developing computational approaches for combinatorial target inference, and supporting training in computational proteomics.

## Background & Summary

Combination therapy is a cornerstone of AML treatment, yet systematically capturing how multiple drugs jointly reconfigure the proteome remains challenging ^1,2^. Traditional synergy metrics quantify abundance-level phenotypic effects of drug pairs but do not reveal underlying protein-level responses or combination-specific mechanisms ^3–5^. To address this, we developed Combinatorial Proteome Integral Solubility/Stability Alteration (CoPISA) assay, a high-throughput proteomics approach that measures protein solubility and stability shifts under single- and multi-drug perturbations. CoPISA provides a readout of protein engagement in lysate, and downstream effects in intact cells, enabling the detection of effects that emerge only under combination treatments.

The dataset encompasses two rational AML drug pairs ^6^ (i.e., LY3009120–sapanisertib (LS) and ruxolitinib–ulixertinib (RU)) applied to four AML cell lines (i.e., NOMO-1, MOLM-13, MOLM-16, and SKM-1) under control, individual drug, and combination conditions in both cell lysate and intact-cell contexts. Protein responses were classified according to whether solubility shifts occur under a single agent, both agents, or exclusively under the combination. This framework operationalizes combination responses using logical-gate concepts ^7^. An “AND-gate” or conjunctional pattern denotes proteins that change significantly only under the combination treatment (A+B) but not under either single agent (A or B), whereas an “OR-gate” or disjunctional pattern denotes proteins responding to at least one single agent and also present in the combination condition (**Fig. 1**). These definitions enable systematic separation of combination-specific and overlapping proteome responses ^8^.

**Figure 1.**
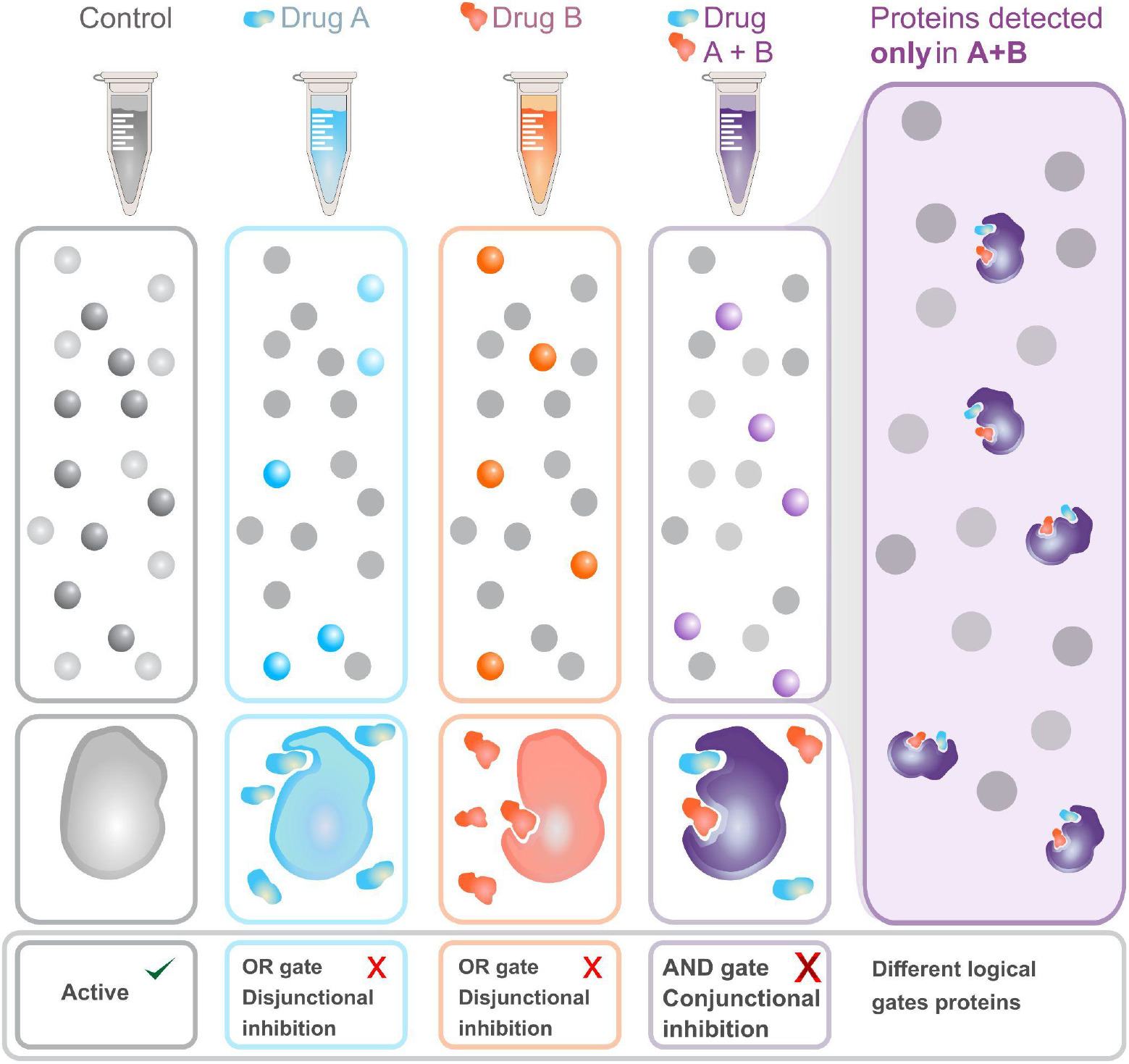
Identification of combination-specific protein responses. Proteomic profiles are compared across three treatment conditions: Drug A, Drug B, and the combination treatment (A+B). Each condition reveals a set of detected proteins, represented as individual protein structures. While some proteins respond to single treatments (A or B), a subset of proteins becomes significantly altered only in the presence of the combination therapy. These proteins are absent or non-significant in both single-agent conditions but emerge under the combined perturbation. The highlighted region illustrates proteins uniquely detected in the A+B condition, representing combination-specific responses consistent with an AND-gate behavior, where a significant effect occurs only when both treatments are present simultaneously.

To interpret these protein responses quantitatively, we classified proteins into logical response categories using statistical significance thresholds applied independently to each treatment contrast. Inspired by Boolean logic used in computer science, we defined two response patterns. An “AND-gate” response refers to proteins that become significantly altered only when both drugs are applied together, while remaining non-significant in either single-agent treatment. This pattern highlights proteins whose response emerges specifically from the combined perturbation, suggesting cooperative or synergistic effects between the drugs. In contrast, an “OR-gate” response refers to proteins that are significantly altered in at least one single-drug treatment and are also detected in the combination condition. These proteins therefore reflect shared or overlapping drug effects rather than combination-specific responses. This logic-based framework enables systematic separation of proteins uniquely driven by drug combinations from those responding to individual drugs (**Fig. 1**).

The resource includes raw and processed multiplexed proteomics data, differential solubility outputs for each treatment and context, intersection analyses, and refined post-translational modification (PTM)-aware identifications. Across four AML cell lines, two assay formats (lysate and intact-cell), four treatment conditions (control, single agent 1 and 2, and combinations), and three technical replicates per condition, the dataset comprises 16 TMT16-plex acquisitions (192 LC–MS/MS raw files). On average, each multiplex experiment yielded several thousands of confidently quantified proteins, providing deep proteome coverage suitable for comparative solubility profiling. The inclusion of pooled reference channels in every TMT set enables within- and cross-plex normalization, and replicate-level concordance supports quantitative reproducibility across biological contexts and treatment modalities. To facilitate automated reprocessing, we provide an SDRF file that encodes sample relationships and experimental metadata ^9, 10^. Data collection is summarized in **Fig. 2**.

**Figure 2.**
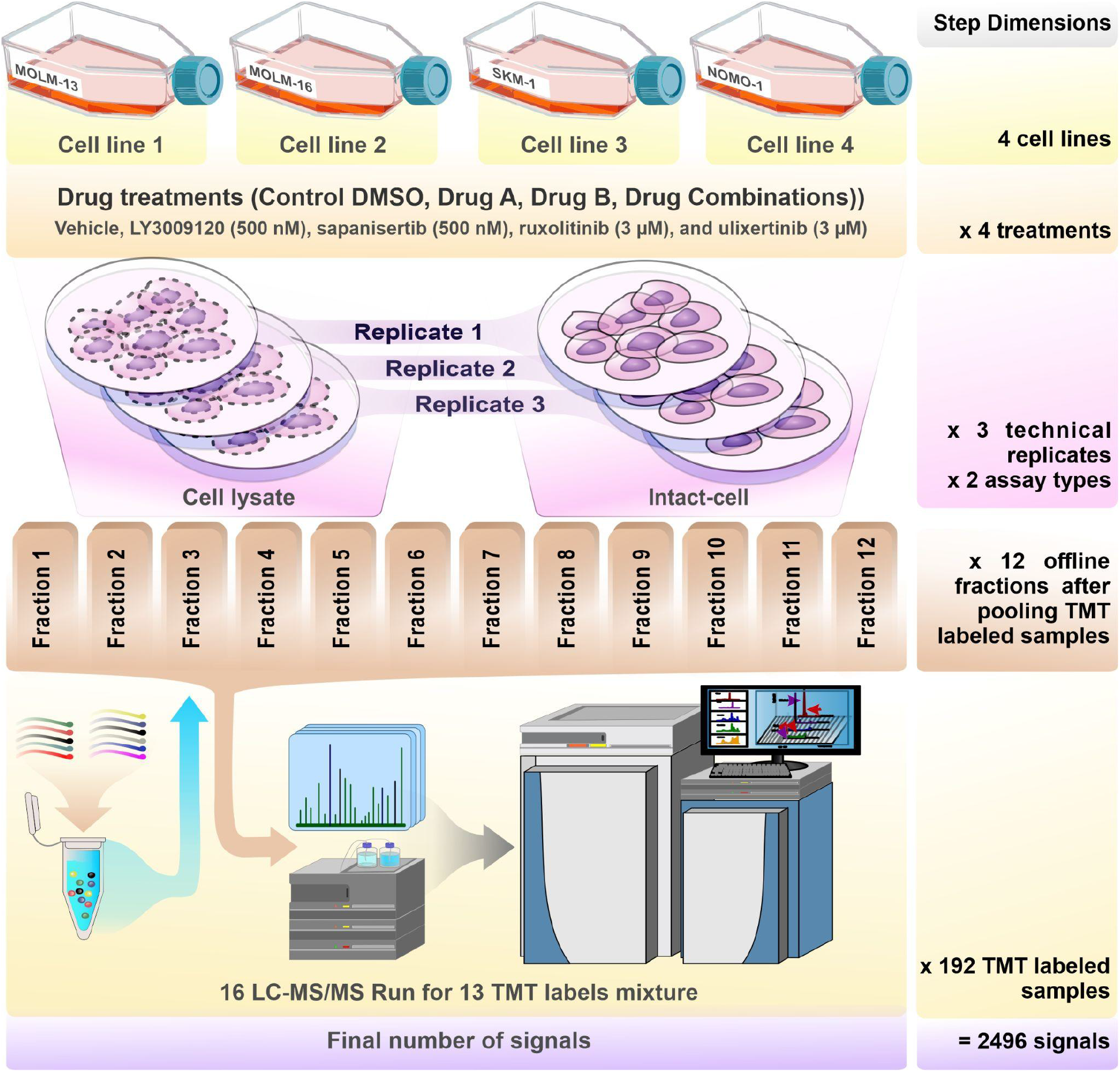
Experimental design and biological and technical replication strategy for CoPISA assay.

These comprehensive data layers support multiple reuse scenarios, including benchmarking computational models for combinatorial target inference, exploring differences between lysate and intact-cell responses, and testing intersection-based analysis strategies for detecting combination-specific protein effects. Cleanly structured and well-annotated perturbation datasets can also serve as reusable inputs for downstream computational frameworks aimed at therapeutic hypothesis generation, including approaches that prioritize candidate vulnerabilities or combination strategies in cancer and other diseases^11^. Furthermore, the open accessibility of both the dataset and the accompanying analysis code makes this resource valuable not only for educational purposes, such as training workshops and teaching modules in computational proteomics, but also for incorporation into public-facing resources and reusable data portals that help other researchers explore, compare, and build upon solubility- and stability-based drug-response datasets^12–15^.

## Methods

### Cell culture

Human AML cell lines MOLM-13, MOLM-16, SKM-1, and NOMO-1 were obtained from the Deutsche Sammlung von Mikroorganismen und Zellkulturen (DSMZ, Germany). Cells were maintained in RPMI-1640 medium (Gibco, Thermo Fisher Scientific, USA) supplemented with 10–20% fetal bovine serum (FBS; Gibco, Thermo Fisher Scientific), 2 mM L-glutamine (Lonza), and 100 U/mL penicillin/streptomycin (Gibco, Thermo Fisher Scientific) at 37 °C in a humidified incubator with 5% CO_2_. Cells were collected by centrifugation at 400 ×g for 4 min. Mycoplasma contamination was excluded using PCR-based testing.

### Drug Treatment Conditions and Thermal Profiling Strategy

To generate protein solubility/stability response profiles, cells or lysates were exposed to defined single agents and combinations at fixed concentrations selected to produce measurable pathway engagement while maintaining compatibility with the CoPISA thermal workflow. The following final concentrations were used: LY3009120 (500 nM), sapanisertib (500 nM), ruxolitinib (3 µM), and ulixertinib (3 µM). Vehicle controls contained 0.1% DMSO. Each sample was distributed across a temperature series spanning 48–59°C in 1°C increments, with a 3 min heat exposure per temperature point and a subsequent 3 min equilibration at room temperature prior to soluble fraction recovery. For each protein, the CoPISA signal per condition was defined as the log2-transformed TMT reporter intensity normalized to the pooled reference channel within each multiplex. Differential solubility (ΔS) was computed as the difference between log2-normalized intensities of treatment and matched DMSO control within the same cell line, assay mode, and multiplex. A “one-pot” pooling strategy was used per treatment condition by combining the soluble material across all temperature points after heating, thereby encoding the area under the melting curve as a single abundance-like measure per protein per condition, enabling high-throughput replication within TMT multiplexing.

### Lysate-based CoPISA sample preparation

For lysate CoPISA, 2 × 10^6^ cells per condition were collected, washed twice in PBS, and resuspended in PBS containing protease inhibitors (Roche). Cells were lysed by four freeze–thaw cycles (liquid nitrogen freeze followed by thaw at 35 °C), and insoluble debris was removed by centrifugation at 10,000 g for 10 min at 4 °C. Clarified lysates were treated with compounds for 15 min at room temperature, split into aliquots for the temperature series, heated, equilibrated, pooled per condition, and ultracentrifuged at 100,000 g for 20 min at 4 °C to isolate the soluble protein fraction used for proteomics.

### Intact-cell CoPISA sample preparation

For intact-cell CoPISA, 2 × 10^6^ cells per condition were treated in culture for 60 min at 37°C and 5% CO_2_, washed twice with PBS, and resuspended in PBS with protease inhibitors. Samples were heated across the same temperature series used for lysates. After equilibration, cells were lysed using four freeze–thaw cycles, pooled per condition, and ultracentrifuged at 100,000 g for 20 min at 4°C to recover soluble proteins. Soluble fractions were snap frozen and stored at −80°C until processing to minimize freeze–thaw variability.

### Sample Preparation and TMT Labeling

Protein concentrations were determined using the Pierce BCA Protein Assay Kit (Thermo Fisher Scientific). For each sample, 27 µg of protein was processed using 10 kDa centrifugal filter units (Amicon Ultra-0.5 mL, 10KD) to ensure consistent buffer exchange and digestion conditions across the dataset. Samples were adjusted with 20 mM EPPS buffer, reduced with 10 mM dithiothreitol (55°C, 30 min), and alkylated with 50 mM iodoacetamide (25°C, 60 min, in the dark). Proteins were digested overnight at 37°C using trypsin (Trypsin/P specificity; enzyme:protein ratio 1:50). Peptides were recovered by centrifugation and additional EPPS rinses.

To support cross-channel comparability within multiplexes, a pooled reference channel was created for each TMT set by combining equal peptide amounts from all samples included in that set. For labeling, 25 µg of peptide per channel was labeled using TMT reagents (Thermo Fisher Scientific) following the manufacturer’s protocol. Labeled channels were combined per TMT set into a single mixture, yielding 325 µg total peptide for each multiplexed experiment (13 channels × 25 µg per channel, including the pooled reference). Across the entire dataset, 16 independent TMT16-plex acquisitions were collected to cover the full matrix of cell lines, assay modes (lysate vs. intact), and treatment conditions.

### Peptide fractionation and LC–MS/MS acquisition parameters

TMT-labeled peptide mixtures were fractionated by high-pH reversed-phase chromatography (Ultimate 3000, Thermo Fisher Scientific) using a gradient from 2% to 15% buffer B over 3 min, to 45% B over 59 min, to 80% B over 3 min with a 9 min hold, followed by re-equilibration to 2% B. Fractions were concatenated to 12 final fractions per TMT set and analyzed independently by LC–MS/MS.

LC–MS/MS was performed on an Orbitrap Eclipse Tribrid mass spectrometer coupled to an Ultimate 3000 nanoLC system with a FAIMS Pro interface (Thermo Fisher Scientific). Peptides were separated on an in-house packed C18 column at nanoflow rates. Data were acquired in data-dependent mode with SPS-MS3 quantification to improve TMT reporter accuracy. MS1 spectra were collected in the Orbitrap, MS2 in the ion trap, and MS3 in the Orbitrap. FAIMS compensation voltages of −40 V and −70 V were alternated. Dynamic exclusion was enabled. Real-time search was performed using a human UniProt database (20,362 entries; download date: 2020-04-17) to facilitate efficient selection and quantification in complex, multiplexed samples.

### Data Records

Raw files were processed in MaxQuant 2.6.5.0 with the Andromeda search engine against the reviewed human Swiss-Prot UniProt database (downloaded 2024-10-10), using a target-decoy strategy and common contaminants. Searches were run in Reporter MS3 mode with manufacturer-provided TMT label correction factors and weighted-ratio normalization to the reference channel. Variable modifications included Met oxidation and protein N-terminal acetylation; Cys carbamidomethylation was fixed. Up to 3 variable modifications and 2 trypsin/P missed cleavages were allowed per peptide. The refined search used the same settings but added PEIMAN2-suggested variable modifications (up to 5 per peptide) ^16^. Precursor mass tolerances were 20 ppm for the first search and 4.5 ppm for the main search, with internal recalibration. Peptides ≥7 amino acids and charges up to +7 were considered. TMT intensities were quantified in MS3 mode with isotope-impurity correction and normalized to the pooled reference. FDR was controlled at 1% at the PSM, peptide, protein, and site levels. Proteins required at least one razor or unique peptide.

The mass spectrometry proteomics data and corresponding metadata have been deposited to the ProteomeXchange Consortium via the PRIDE partner repository with the dataset identifier PXD066812. The raw data is organized in 16 archived files, corresponding to 16 TMT-labelled samples, and named as Kit *i*_*j*, where *i* is the TMT-labelling kit number, and *j* is the ordinal number of the sample that was labelled with Kit *i*. Every archive file contains 12 raw mass spectrometry data, corresponding to 12 fractions obtained from 16 samples. In total there are 192 raw files. Moreover, raw files were converted to the HUPO proteomics standard mzML. Unprocessed MaxQuant searches were deposited as 2 archived files. To allow the community to reproduce the MaxQuant search, we also deposited parameter files for the search software, as well as the protein database that was used in our search. Lastly, the SDRF file was also published in order to provide transparency on the experimental design and the raw files ^14^.

### Technical Validation

To support reuse of this dataset for independent reanalysis and method development, we performed technical validation at three levels: (i) biological input quality, (ii) experimental reproducibility of the CoPISA perturbation workflow in both extract and intact formats, and (iii) analytical performance of TMT-based multiplexed LC–MS/MS. The final processed dataset comprised 42,083 filtered peptides, which were aggregated to 5,559 proteins across 191 samples, providing broad proteome coverage and enabling proteome-wide assessment of drug-induced solubility changes across diverse cellular pathways. Quantitative precision across biological replicates was assessed using the coefficient of variation (CV) of protein intensities within experimental conditions. Median protein CV across replicate groups was 12.32% (interquartile range, 10.91–15.61%), supporting the robustness of the multiplexed quantitative measurements.

All validation steps were designed to identify and remove problematic runs, confirm consistency of multiplexed quantification, and demonstrate that combination treatments generate non-trivial proteome-wide solubility signatures rather than artifacts driven by acquisition or processing variability. All AML cell lines (MOLM-13, MOLM-16, SKM-1, NOMO-1) were obtained from a major biological resource center (DSMZ) and maintained under standardized culture conditions to minimize variability attributable to media composition, density, or handling.

CoPISA profiling was performed in two complementary modes, lysate-based (extract) and living-cell (intact), using the same drug concentrations, thermal challenge design, and downstream protein isolation strategy. The dual-format design provides an internal validation axis: extract emphasizes direct physicochemical effects consistent with primary drug–protein engagement in lysates, whereas intact captures integrated cellular responses that include downstream remodeling ^17,18^. For each cell line and condition, three technical replicates were acquired, enabling replicate-level concordance assessment within the same biological background. Temperature challenges were executed using a fixed program (3 min heating followed by 3 min equilibration) across 48–59°C in 1°C increments, and solubility readouts were derived from identically pooled temperature-series material per treatment. The use of identical heating grids, pooling strategy, and ultracentrifugation conditions across all samples constrains technical degrees of freedom and supports comparability across treatments, cell lines, and preparation modes.

In addition to replicate-aware experimental design, run-level quality control was assessed directly from reporter ion intensity distributions across TMT channels and multiplex sets. As shown in **Fig. 3**, log2-transformed reporter intensity densities and boxplots were broadly concordant across TMT labels, runs, and multiplexes, indicating stable signal behavior and absence of systematic channel- or run-specific distortion. These visualizations complement replicate-level reproducibility, confirming that the multiplexed acquisition was technically consistent across the dataset before downstream normalization and protein-level inference.

**Figure 3.**
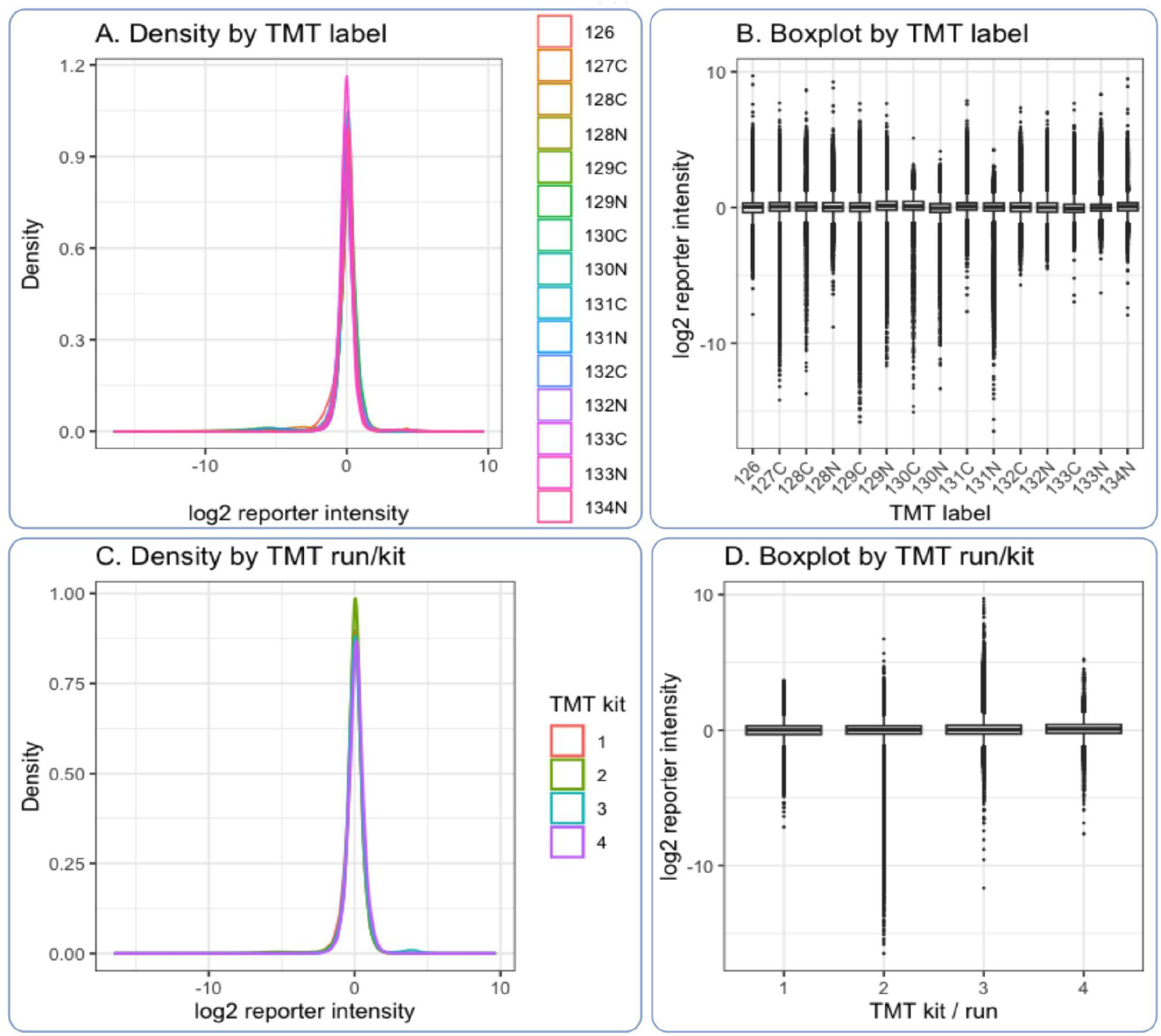
Quality-control assessment across TMT labels and multiplex runs. (A) Density distributions of log2-transformed reporter ion intensities stratified by TMT label. (B) Boxplots of log2-transformed reporter ion intensities across TMT labels. (C) Density distributions of log2-transformed reporter ion intensities stratified by TMT run/kit. (D) Boxplots of log2-transformed reporter ion intensities across TMT runs/kits.

To control variability introduced during proteomic processing, total protein input was standardized by BCA quantification before digestion, and equal peptide amounts were used for TMT labeling. Sample processing employed established reduction/alkylation conditions and filter-aided cleanup before overnight trypsin digestion, which reduces interference from detergents and low-molecular contaminants and improves digestion reproducibility. TMT labeling was performed using consistent peptide input per channel, and each multiplex set included a pooled reference channel generated from equal peptide amounts across all samples in that set. The reference channel supports within-plex normalization and enables cross-sample comparison by anchoring reporter intensities to a common internal standard. Reporter ion intensities were normalized in two stages. First, within-plex normalization was performed by dividing each channel by the pooled reference channel to correct for loading differences. Second, between-plex normalization was applied using median centering of protein-level log2 ratios across multiplexes to minimize batch-specific shifts while preserving biological contrasts. Fractionation by high-pH reversed-phase chromatography was performed using a fixed gradient and concatenation scheme to yield 12 fractions per multiplex set, increasing proteome coverage and reducing ratio compression by distributing precursor complexity across runs.

To further document analytical consistency after preprocessing, we summarized sample-level normalized intensity profiles and protein-level coverage across retained channels. After filtering reverse hits, contaminants, pooled or excluded channels, features with insufficient peptide support, and non-quantified entries, peptide-level measurements were median-centered before aggregation. The resulting protein-level matrix preserved broad sample comparability while maintaining biologically meaningful treatment contrasts, supporting its suitability for downstream differential solubility modeling and method benchmarking.

All TMT fractions were analyzed on the same LC–MS/MS platform (Orbitrap Eclipse Tribrid with FAIMS Pro), using SPS-MS3 acquisition to improve quantitative accuracy of reporter ions in multiplexed experiments. Alternating FAIMS compensation voltages (−40 V and −70 V) were used to enhance depth and reduce chemical noise. Because each TMT set contains an internal pooled reference channel, channel-wise reporter distributions can be inspected for gross outliers indicative of labeling failures, injection anomalies, or fraction-level losses. Statistical inference of differential solubility was performed at the protein level using linear modeling across replicate-level log2-normalized reporter intensities. For each cell line and assay mode, contrasts were defined comparing single agents and combinations to their matched DMSO controls. Empirical Bayes moderation was applied to variance estimates to improve stability across the proteome. In addition, several orthogonal QC summaries were used to evaluate quantitative consistency, including channel-level intensity distributions, run/kit-level signal concordance, feature missingness as a function of abundance, per-sample protein coverage, and sample-to-sample correlation structure (**Fig. 3**). The repeated appearance of technical replicates within the same multiplex set further enables detection of run- or fraction-specific deviations.

## Usage Notes

The goal of this work was to identify protein targets of novel combination therapy in AML. The PRIDE project PXD066812 is the only source. The researchers can reuse the data in different ways. The unprocessed RAW files can be downloaded from the PRIDE and searched by researchers with the proteomics database search software of their choice. In this case, only RAW files and SDRF are needed to set the experimental design inside the proteomics software. The users can use them directly for preprocessing and downstream analysis. The preprocessing and downstream analysis for MaxQuant results can be found in the Code Availability section.

In case researchers want to reproduce our search, we provided parameter files for the software mentioned above. Researchers need to download the.RAW files, parameters for the software, and the proteomics database (FASTA file), and install the proteomics software. The paths inside the parameter file should be adjusted to the appropriate paths on the local machine from which the analysis is run. As some software for proteomics requires mzML files, we also provided those to reduce the time for analysis. Additionally, if researchers would like to search the data with slightly different parameters, they can change them in the parameter file, save it under a new name, and run the search.

## Data Availability

The mass spectrometry proteomics data and corresponding metadata are available at the ProteomeXchange Consortium via the PRIDE partner repository with the dataset identifier PXD066812 (https://www.ebi.ac.uk/pride/archive/projects/PXD066812).

## Code Availability

The R code used for reporter normalization, protein-level aggregation, statistical modeling, logical gate classification (AND/OR patterns), and visualization is available at the GitLab repository (https://version.helsinki.fi/ehzange/Max_RULS). The repository includes reproducible scripts for data import, quality control diagnostics, normalization workflows, linear modeling, multiple testing correction, and generation of intersectional solubility maps. All analyses were performed using R (version 4.5.2) with documented package dependencies to ensure computational reproducibility.

## Author Contributions

E.Z. led manuscript writing, computational analyses, data interpretation, data visualizations, and scientific illustrations, while E.G. led the experimental analyses and contributed to data interpretation, U.V. participated in proteomics data analysis, and D.R. conducted the proteomics mass spectrometry experiments and data analysis. M.J. conceived the study and designed the research questions, also provided scientific direction, and supervised the project. All authors contributed to the manuscript preparation.

## References

1. Cherry, E. M. et al. Venetoclax and azacitidine compared with induction chemotherapy for newly diagnosed patients with acute myeloid leukemia. Blood Adv. 5, 5565–5573 (2021).

2. Barneh, F. et al. Integrated use of bioinformatic resources reveals that co-targeting of histone deacetylases, IKBK and SRC inhibits epithelial-mesenchymal transition in cancer. Brief Bioinform 20, 717–731 (2019).

3. Tyner, J. et al. Functional genomic landscape of acute myeloid leukemia. Nature 562, 526–531 (2018).

4. Mirzaie, M. et al. Designing patient-oriented combination therapies for acute myeloid leukemia based on efficacy/toxicity integration and bipartite network modeling. Oncogenesis 13, 11 (2024).

5. Barneh, F. et al. Valproic acid inhibits the protective effects of stromal cells against chemotherapy in breast cancer: Insights from proteomics and systems biology. J Cell Biochem 119, 9270–9283 (2018).

6. Targeting acute myeloid leukemia resistance with two novel combinations demonstrate superior efficacy in TP53, HLA-B, MUC4 and FLT3 mutations. Biomedicine & Pharmacotherapy 192, 118647 (2025).

7. Jafari, M., Ansari-Pour, N., Azimzadeh, S. & Mirzaie, M. A logic-based dynamic modeling approach to explicate the evolution of the central dogma of molecular biology. PLoS One 12, e0189922 (2017).

8. Gholizadeh, E. et al. Shifting beyond classical drug synergy in combinatorial therapy through solubility alterations. bioRxiv (2024) doi:10.1101/2024.11.08.618644.

9. Dai, C. et al. A proteomics sample metadata representation for multiomics integration and big data analysis. Nat. Commun. 12, 5854 (2021).

10. Claeys, T. et al. lesSDRF is more: maximizing the value of proteomics data through streamlined metadata annotation. Nat. Commun. 14, 6743 (2023).

11. Zangene, E.Marashi, S.-A. & Montazeri, H. SL-scan identifies synthetic lethal interactions in cancer using metabolic networks. Sci Rep 13, 15763 (2023).

12. Webel, H., Perez-Riverol, Y., Nielsen, A. B. & Rasmussen, S. Mass spectrometry-based proteomics data from thousands of HeLa control samples. Scientific Data 11, 112 (2024).

13. Ping, L. et al. Global quantitative analysis of the human brain proteome and phosphoproteome in Alzheimer’s disease. Scientific Data 7, 315 (2020).

14. Vadadokhau, U. et al. Preventing Proteomics Data Tombs Through Collective Responsibility and Community Engagement. Sci Data 13, (2026).

15. Zangene, E. et al. DORSSAA: Drug-target interactOmics Resource based on Stability/Solubility Alteration Assay. bioRxiv (2023) doi:10.1101/2023.12.29.573639.

16. Nickchi, P. et al. Monitoring Functional Posttranslational Modifications Using a Data-Driven Proteome Informatic Pipeline. Proteomics 25, e202400238 (2025).

17. Martinez Molina, D. et al. Monitoring drug target engagement in cells and tissues using the cellular thermal shift assay. Science 341, 84–87 (2013).

18. Gaetani, M. et al. Proteome Integral Solubility Alteration: A High-Throughput Proteomics Assay for Target Deconvolution. J Proteome Res 18, 4027–4037 (2019).

